# Light-harvesting strategies and competition drive niche partitioning among *Ostreobium* lineages in the spectral architecture of the coral reef

**DOI:** 10.64898/2026.03.02.709049

**Authors:** Marisa M. Pasella, Manuel Poretti, Alasdair Sim, Francesco Ricci, Felix Powrie, Heroen Verbruggen

## Abstract

*Ostreobium*, a siphonous green alga capable of living inside of calcium carbonate substrates, including the skeletons of reef-building corals. This study investigates spectral niche preferences and physiological strategies of *Ostreobium* using community-wide experiments. We exposed natural *Ostreobium* communities from *Porites lutea* collected across shallow, mid, and deeper-water sites to three light conditions: far-red, blue, and white light, simulating healthy shallow-water corals, deeper water conditions, and bleached coral skeletons respectively. Using 16S rRNA metabarcoding and chlorophyll analysis, we assessed community changes and physiological responses over 16 weeks. We show significant variation in spectral preferences among *Ostreobium* OTUs, with clear evidence for both generalist and specialist strategies. Chlorophyll analysis showed photoacclimation responses through changes in pigment compositions. Our work shows that the spectral architecture of the reef plays a role in structuring Ostreobium communities, but the many mismatches between spectral preferences of OTUs and their observed presence in nature, suggests that inter-species competition is likely to be an even stronger contributor to community structure across the reef’s microhabitats. We show that physiological heterogeneity within *Ostreobium* is strongly phylogenetically structured, and our results clearly highlight the importance of considering OTU-level differences when predicting community responses to environmental disturbances such as coral bleaching. While generalist OTUs dominate natural communities, these do poorly in incubations, and we hypothesise that white light specialists may become key players during coral bleaching events. Our work is a substantial advance in our understanding of *Ostreobium* ecology and provides a framework for interpreting future environmental sequencing data, offering insights into the functional roles of the different OTUs.

## Introduction

*Ostreobium* is a siphonous green alga capable of dissolving calcium carbonate, capable of burrowing through limestone rock by dissolving calcium carbonate (Tribollet et al., 2019; Verbruggen & Tribollet, 2011). It colonises and survives in a range of temperate to tropical habitats including bivalve shells and in the skeletons of living corals (Tandon, Pasella, et al., 2023), and has been found at depths of up to 200m (Aponte & Ballantine, 2001; Littler et al., 1985).

Inside the coral skeleton, *Ostreobium* is diverse, with at least ca. 80 operational taxonomic units (OTUs) at or near the species-level organised into four major lineages and several distinctive clades (del Campo et al., 2017; Giraldo-Vaca & Sánchez, 2024; Gutner-Hoch & Fine, 2011; Marcelino & Verbruggen, 2016). Inside the skeleton, they carry out oxygenic photosynthesis as part of a more complex microbiota (Ricci et al., 2019, 2023; Tandon, Ricci, et al., 2023). How the niche features of the coral skeleton may impact the diversity and distribution of *Ostreobium* has not yet been investigated in a systematic way, but several studies can help us form hypotheses.

Caribbean *Ostreobium* communities did not correlate with nutrient load associated with river runoff (Gonzalez-Zapata et al., 2018), but genotypic variation was observed based on the depth of collection of the coral host. Similarly, *Ostreobium* distributions correlated with depth in *Porites lutea* and *Goniastrea perisi* corals in the Red Sea (Gutner-Hoch & Fine, 2011). The biomass of *Ostreobium* is also known to correlate well with skeletal morphology, with corals having larger calice structures being more effective in delivering light to the skeleton (Fordyce et al., 2021). Both these observations contribute to the notion that light may affect *Ostreobium* distribution and diversity.

The coral reef presents a complex framework of microhabitats with different light intensities and wavelenghts of light, and the architecture of this spectral diversity of light can be expected to influence the distribution of organisms that depend on light for their energy provision, *Ostreobium* included. Irradiances reaching the coral skeleton can vary considerably in intensity and spectrum (Magnusson et al., 2007). Symbiodiniaceae in the tissue absorb most photosynthetically active radiation (PAR), and only an estimated 0.1%–1% of light reaching the coral tissue is transmitted into the skeleton (Halldal, 1968; Kanwisher & Wainwright, 1967; Schlichter et al., 2008). Blue light is generally extremely depleted in the skeletal light environment (Magnusson et al., 2007). However, low-energy far-red radiation (> 700nm) is not used much by Symbiodiniaceae, and between 56% and 145% of the incidental light of these wavelengths are estimated to reach the *Ostreobium* layer in the skeleton, with the higher-than-100% values likely explained by the light-scattering properties of the skeleton (Magnusson et al., 2007).

At least some *Ostreobium* strains can perform photosynthesis using far-red wavelengths (Wilhelm & Jakob, 2006), which may explain its abundance in the skeleton of shallow-water corals. However, far-red photosynthesis is no longer advantageous in water deeper than 5-10m as far-red wavelengths are attenuated in the water column, leaving any *Ostreobium* growing in deeper water in very low light, mostly consisting of the blue wavelengths. This begs the question whether *Ostreobium* strains can acclimate to those very divergent light environments or whether this is due to niche differentiation, with specialisation of different *Ostreobium* strains to particular light environments.

In healthy corals, the skeleton may often experience low levels of PAR irradiance, but this is quite different during coral bleaching events. With the Symbiodiniaceae largely gone from the coral tissue during such events, nearly no light is absorbed in the tissue layer, and much more light, likely including more PAR wavelengths, reaches *Ostreobium* in the skeleton. This gives rise to a puzzling contradiction, because even though *Ostreobium* is considered a low-light specialist, it is often observed to bloom when corals bleach (Fine et al., 2005; Fordyce et al., 2021). One would expect such intense light to act as a stressor and influence the *Ostreobium* community, since some studies demonstrated photosynthesis saturation at relatively low light levels (Galindo-Martínez et al., 2022; Ralph et al., 2007).

These observations show that *Ostreobium* can thrive across a broad spectrum of lightscapes, yet it remains unclear whether this is due to strong acclimation potential of *Ostreobium* strains or spectral niche differentiation and specialisation among lineages. Based on the observed species diversity in the genus, one certainly could argue that there is potential for specialisation. This notion is supported by the diverse absorption and fluorescence spectra between *Ostreobium* strains (Koehne et al., 1999), with the primary distinction lying in the amount of chlorophyll *b*. The notion that species are structured across depth gradients further suggests light may be an important structuring component for Ostreobium communities (Gonzalez-Zapata et al., 2018; Gutner-Hoch & Fine, 2011). Despite these insights, our understanding of the spectral niche of *Ostreobium* remains fragmentary.

The goal of this study is to investigate spectral specialisation among *Ostreobium* strains using community-wide experiments. Specifically, we aim to (1) test the hypothesis that light spectrum influences the composition of *Ostreobium* communities, (2) evaluate changes in pigment composition as a function of light spectrum, and (3) gain understanding of the distribution of photobiology-relevant traits across the diversity of *Ostreobium* OTUs. We approach these questions with experimental exposure of natural *Ostreobium* communities to a range of relevant light conditions: (1) far-red, simulating the skeleton of healthy shallow-water corals, (2) blue, simulating deeper water conditions and (3) white, simulating the skeleton of bleached corals. We use metabarcoding to document community changes and monitor the change in chlorophylls.

## Methods

### Sample collection and processing

Fragments of thirty-eight colonies of *Porites lutea* were collected from Heron Island (Queensland, Great Barrier Reef) in January 2020 (Fig. 1 and Supplementary Figure S1). Eighteen samples were collected at water depth <1m, from the research zone of the Reef flat (23°44’S, 151°91’E). Twelve samples were collected between 3m and 10m depth in the Tenements and Canyons sites (respectively 23°25’S, 151°55’E and 23°27’ S, 151°55’E), and eight samples were collected at water depth > 15m (Libby’s site: 23°26’S, 151°57’E). Samples were collected using hammer and chisel, and placed in new zip-lock polyethylene bags in seawater. We collected from the top of the coral colonies to avoid sampling the more shaded lateral parts. We cut out the visible green layer of *Ostreobium* from each skeleton and divided it into eight subsamples (∼ 0.3cm3 each). One subsample was flash-frozen in liquid nitrogen and kept at -80° for 16S rRNA metabarcoding. One was incubated in 100% methanol for chlorophyll assessment. The remaining six subsamples were used in a sixteen-week experimental incubation.

**Figure 1.**
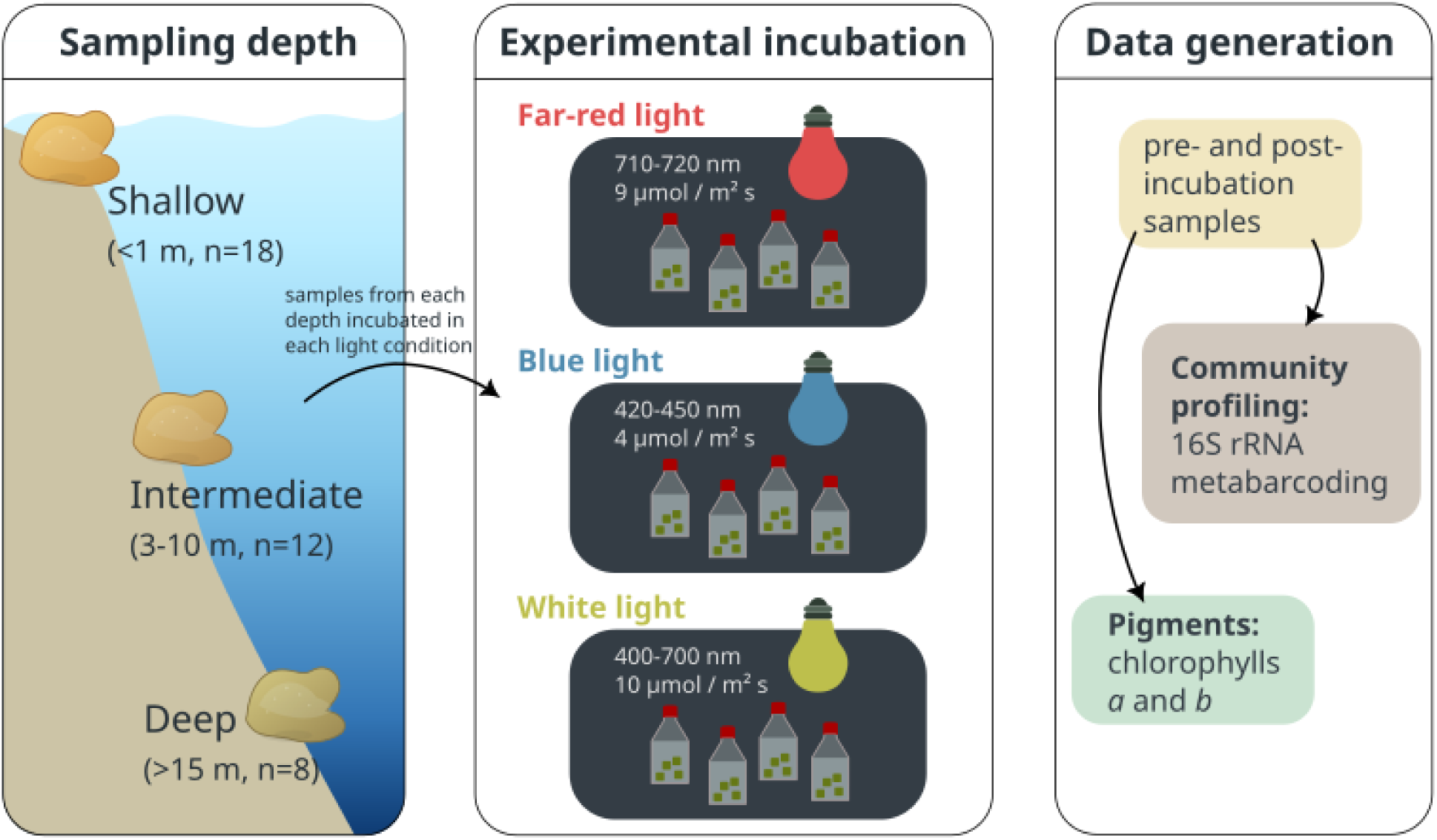
Study design. Coral fragments collected from different depths were incubated in 3 different light conditions: far-red, blue and white. Metabarcoding of 16S rRNA and chla and chlb quantification was performed before and after incubation.

### Experimental setup

To evaluate changes in chlorophyll concentrations and *Ostreobium* community composition in different light spectra, we incubated the *Ostreobium*-containing skeleton fragments in different light conditions (Fig. 1). The three light conditions were: (1) far-red light (710-720m) at 9 µmol m-2s-1 photons, (2) blue light (420-450nm) at 4 µmol m-2s-1 photons and (3) white cool light (400-700nm) at 10 µmol m-2s-1 photons.

For the white light spectrum, we based our light intensity setup on the estimated maximum midday light intensity reaching coral tissue, approximately 2000 µmol photons m⁻² s⁻¹ (Veal et al., 2010). Previous studies suggest that only about 0.1% of this incident radiation penetrates the coral tissue to reach the skeleton (Kanwisher and Wainwright, 1967; Halldal, 1968; Schlichter et al., 1997). Consequently, the *Ostreobium* layer within the coral skeleton is exposed to approximately 1-2 µmol photons m⁻² s⁻¹ of the full light spectrum. To simulate an increase in light intensity within the coral skeleton (as would be the case during coral bleaching), we elevated this value by a factor of 5-10, setting the light intensity to 10 µmol photons m⁻² s⁻¹.

For the blue light setup, we applied a lower light intensity of 4 µmol photons m⁻² s⁻¹ to reflect the lower-intensity blue light in deeper waters. For the far-red light setup, we provided a light intensity comparable to the white light setup, at approximately 9 µmol photons m⁻² s⁻¹. This choice is based on published findings that zooxanthellae in coral tissue are largely unable to utilize far-red light. Consequently, far-red light penetrates more effectively to the *Ostreobium* layer within the coral skeleton (Magnusson et al., 2007; Shibata & Haxo, 1969).

The incubated fragments were maintained in 10 ml f/2 medium in a 25 ml tissue culture flask, at 26°C on a 12h:12h light:dark cycle. The set-up using black plastic boxes with custom lighting is shown in Fig. S2. The light:dark cycle was offset from the external light-dark cycle to facilitate carrying out media changes at night, using only the light from within the boxes, thereby avoiding exposure of the samples to wavelengths of light other than those of the experimental treatment. At the end of the incubation period, we froze part of the samples for 16S rRNA metabarcoding and kept others in 100% methanol for pigment extractions.

### Chlorophyll analysis

Chlorophyll *a* (chl*a*) and chlorophyll *b* (chl*b*) were measured (Porra et al., 1989) and expressed in µg ml⁻¹, with the volume referring to the solvent used. The difference in chlorophyll concentrations between the different light conditions were assessed using 1-way ANOVA (n = 72 for shallow-water samples, n = 48 for samples from intermediate depths, and n = 32 for deeper-water samples). When significant differences for the main effect were observed (p < 0.05), a Tukey’s pairwise comparison test was performed. The same protocol was applied for differences between the chl*b*/chl*a* ratio.

### DNA extraction and metabarcoding

DNA was extracted from snap-frozen skeletal sub-samples using the Wizard Genomic DNA Purification Kit (Promega). We used a two-step PCR protocol to produce a multiplexed library of chloroplast 16S rRNA genes. The first PCR used 16S-specific primers with universal adapter overhangs, and the second reaction attached Illumina adapters containing unique library indices for high-throughput sequencing.

A new set of primers was designed to target the combined v4 and v5 variable regions of algal 16S rRNA genes. Based on an alignment of 229 taxonomically diverse algal chloroplast 16S rRNA genes (including *Ostreobium*), we identified the flanking sites of these variable regions, which were scanned manually for regions of high mean pairwise identity that could serve as primer binding sites. To obtain the primers used for our first PCR reaction, we added universal adapter overhangs to these 16S-specific sequences, leading to these primer sequences: v3-v4F: 5’-<u>GTGACCTATGAACTCAGGAGTC</u>GCCAGCMGCCGCGGTAA − 3′, and v5-v6R: 5′-<u>CTGAGACTTGCACATCGCAGC</u>ATCGAATTAAACCACATGCTCCAC − 3′, where the underlined parts are the universal overhangs. The first round of PCR was conducted in 25 µL reactions using NEB TAQ DNA polymerase, 10 pmol of each primer and 1 µL of diluted template DNA, with the thermal cycling protocol consisting of initial denaturation (30s at 95°C), 18 cycles of 20s at 95°, 30s at 58°, 30s at 68°, and a final extension of 5m at 68°. Products of the first PCR were cleaned using CleanNGS magnetic beads at a 1:1 ratio.

The second round of PCR was used to attach Illumina adapters with sample-specific indices to the products of the previous PCR round (Ricci et al., 2023). This was conducted in 20 µL reactions using the GoTaq Greenmix and 5 pmol of unique forward and reverse barcoded Illumina indexes. The thermal cycling protocol consisted of initial denaturation (3m at 95°C), 15 cycles of 15s at 95°, 30s at 60°, 30s at 72°, and a final extension of 7m at 72°.

A combined library was prepared by pooling 5 µL from each well and performing another cleanup step using a 4:5 ratio of pooled DNA to CleanNGS beads. This solution was used for an Illumina Nextseq 550 sequencing run at the Walter and Eliza Hall Institute of Medical Research in Melbourne (Australia).

### Analysis of 16S rRNA metabarcodes and phylogeny

Demultiplexed sequencing data was imported into QIIME2 v2022.11.1 (Kuczynski et al., 2011) and sequencing adapters (forward: GTGACCTATGAACTCAGGAGTC, reverse: CTGAGACTTGCACATCGCAGC) were trimmed using “qiime cutadapt trim-paired”. A summary of the trimmed data was generated with “qiime demux summarize” to identify regions with low sequencing quality to be removed. The next steps were performed with the DADA2 pipeline v1.28.0 (Callahan et al., 2016) following the guidelines at https://iallali.github.io/DADA2_pipeline. Trimmed data was filtered using parameters maxN=0, truncQ=2, rm.phix=TRUE and maxEE=2. The error model was calculated and used to run the core sample inference algorithm on forward and reverse reads. Paired reads were merged and the amplicon sequence variant (ASV) table generated. Finally, chimeric ASVs were first eliminated *de novo* using the R function removeBimeraDenovo and then using the reference-based VSEARCH uchime_ref method (Rognes et al., 2016) against the non-redundant small subunit (SSU) SILVA database (release 138).

ASVs were filtered to retain only green algae using BLASTn against available *Ostreobium* 16S rRNA (GenBank accessions: OK189523-31; PP315243-78; MH894197, MH894198, KU979012). We settled on a percentage identity threshold of 87, which based on our preliminary experimentation retained a wide range of *Ostreobium* strains along with some off-target algae. Further filtering of sequences to retain only those in the Ulvophyceae *Ostreobium* clade was done based on blastn searches of the ASV sequences against the Genbank nt database and preliminary phylogenetic trees.

Operational taxonomic units (OTUs) were defined by clustering ASVs with at least 98% sequence homology. This was done using the R function Treeline (from package DECIPHER) with parameters method=complete, cutoff=0.02 and type=clusters. Low-abundance OTUs (< 100 reads and < 3 samples) were removed. OTU abundance (number of reads) was normalized across samples using a variance stabilizing transformation with the R functions estimateSizeFactors(type = “poscounts”) and varianceStabilizingTransformation from the R package DESeq2 v1.40.2. Next, the R function ordinate from the Phyloseq v1.50.0 package was used to perform an ordination of the normalized number of reads with method RDA. Finally, the function scores from Vegan v2.6-8 was used to extract the scores (site and species coordinates) that were used to draw the biplots using ggplot2 v3.5.1. Boxplots were generated to compare the abundance of the most prevalent *Ostreobium* OTUs throughout different light treatments. Separate graphs were made for samples originating from different depths.

To determine the phylogenetic relationships among OTUs and identify their relationships to known green algal lineages, we built a phylogeny from an alignment of 220 plastid 16S rRNA sequences. These include the OTUs obtained here, 16S sequences from chloroplast genomes of the Bryopsidales and other Ulvophyceae orders (Cremen et al., 2019; Marcelino et al., 2016; Pasella et al., 2022; Verbruggen et al., 2017), sequences of previously isolated culture strains (Sauvage et al., 2016) available on Genbank, 16S sequences from the PR2 v5.1.1 database (Guillou et al., 2013), and the environmental *Ostreobium* sequences from del Campo et al. (2017). To help reduce the total number of sequences, we clustered some datasets using the --cluster_fast function in VSEARCH v2.18.0 (Rognes et al., 2016). This includes the datasets from del Campo (at 0.94 identity), Sauvage strains (at 0.98 identity) and PR2 (at 0.96 identity). Sequences were aligned using MAFFT v7.505 (Katoh & Standley, 2013) with default settings and automatic determination of sequence direction. The phylogenetic tree was inferred with maximum likelihood using IQ-Tree v2.1.4 (Minh et al., 2020), with the GTR+F+I+G4 as the model determined by the built-in ModelFinder and 1,000 ultrafast bootstrap replicates to estimate branch support (Hoang et al., 2018; Kalyaanamoorthy et al., 2017).

### Evaluating specialisation

To analyse the response of OTUs to particular light incubations we calculated the log-fold change (LFC) in OTU abundance between the pre-treatment sample and the samples incubated in different light spectra like this example for blue light:

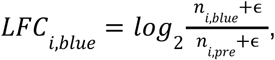

where *i* is the OTU, *n_i,blue_*and *n_i,pre_* are the normalised abundances in the blue treatment and pre-treatment samples, respectively, and ɛ = 0.001 as a pseudocount to add to handle zeros in log calculations. These values were averaged across all samples to obtain a mean LFC (and standard error) for each OTU and light condition.

This was done to achieve two things. First, we wanted to evaluate whether OTUs are specialists in particular light conditions or generalists. To this goal, we calculated a selection index (SI) as follows:

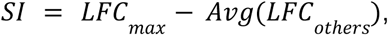

where *LFC_max_* is the LFC of the light incubation that had the highest LFC, and *Avg(LFC_others_)* is the average LFC of the remaining two light incubations. We used these values to classify OTUs as generalists (low selectivity, *SI* < 3.16) or specialists (strong selectivity, *SI* ≥ 3.16). This means, for instance, that for an OTU to qualify as a blue light specialist, its abundance after incubation in the blue light treatment has be 10-fold (2^3.16^) higher than the average abundance in the far-red and white light treatments (averaged across all samples).

Second, we wanted to evaluate which OTUs showed good performance growing in far red light, which was done based on its performance in far-red light *LFC_far-red_*using these thresholds:

- Thrives: *LFC_far-red_* ≥ 0 (increase in relative abundance)
- Intermediate: -3.16 ≤ *LFC_far-red_* < 0 (abundance drop by less than 10x)
- Poor: *LFC_far-red_* < -3.16 (abundance drops by more than 10x)

## Results

### Substantial *Ostreobium* biodiversity is found in GBR *Porites* corals

Analysis of the 16S rRNA metabarcodes revealed a total of 39,295 ASVs before any filtering. BLAST searches indicated that 246 ASVs may belong to the *Ostreobium* clade, and these ASVs clustered into 53 OTUs at the 98% level. Further taxonomic filtering resulted in the final set of 39 OTUs belonging to the Ulvophyceae green algae. The median length of the OTU sequences was 449 nt.

The phylogeny showed the various lineages of Ulvophyceae used in the dataset (Fig. 2a), and the great majority of OTUs obtained from the coral skeleton samples fell in the *Ostreobium* clade (suborder Ostreobineae of the order Bryopsidales). Several lineages could be clearly recognised within the *Ostreobium* clade, indicated to the right of Fig. 2A using the clade assignments that were used in previous papers (del Campo et al., 2017; Marcelino & Verbruggen, 2016; Sauvage et al., 2016). In addition to the four lineages (L1-4) recognised by Marcelino & Verbruggen (2016), which we will use as our main reference for classification, a new deep-branching *Ostreobium* OTU (101) was recovered from our study, which we will refer to as lineage 5 (L5). A few OTUs fell outside the Ostreobineae suborder, instead representing likely endolithic stages of species from the Bryopsidineae (OTUs 32, 54, 99) and Halimedineae (OTUs 33). The Halimedineae also include two endolithic lineages, likely represented here by OTUs 3, 56 and 56. Two OTUs fell in non-focal groups including the Ulvales-Ulotrichales lineage (OTU 109) and an unknown deep-branching Ulvophyceae (OTU 7).

**Figure 2.**
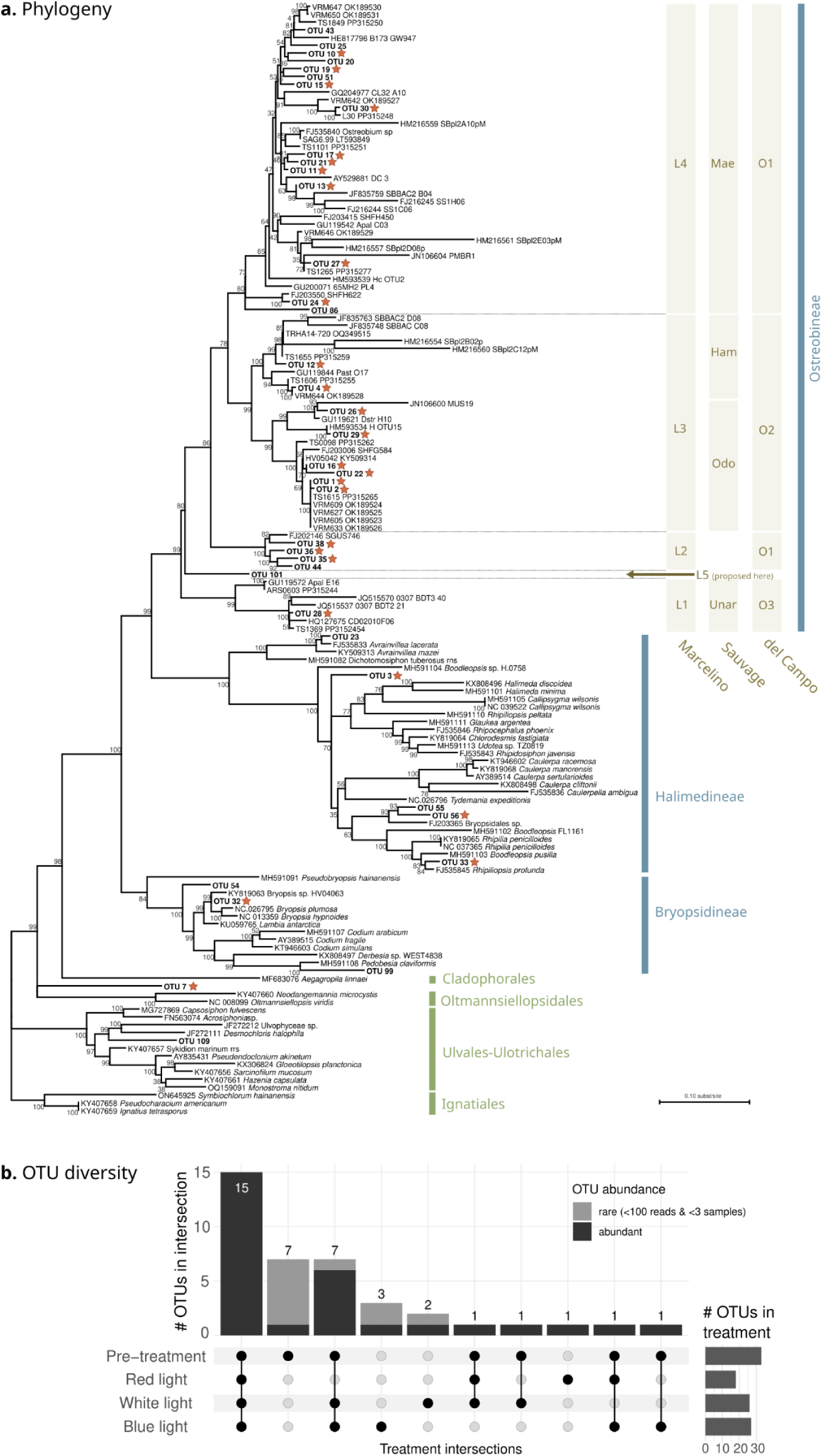
*Ostreobium* diversity and distribution. **(a)** Phylogeny of the 16S rRNA gene showing the position of the OTUs from this study (boldface) with culture strains and formerly reported environmental sequences. The three *Ostreobineae* classification frameworks are indicated to the right (see text), and red stars indicate common OTUs. **(b)** Upset plot showing the abundance of *Ostreobium* OTU diversity across light treatments, regardless of sample depth. See Suppl. Fig. S3 for depth-specific upset plots.

Upset plots illustrating the presence of OTUs across pre-treatment samples (i.e. samples collected from nature before being subjected to treatment) and samples deriving from our light treatments showed that the community is characterized by a combination of rare and abundant OTUs (Fig. 2b). Many rare OTUs were only found in pre-treatment samples. The highly abundant OTUs were often present in multiple light treatments. Indeed, the 32 OTUs detected following light incubation were enriched with highly abundant OTUs (∼91%), most of which were seen in multiple light treatments, while OTUs specific to particular light incubation types were often rare. Only 3, 2 and 1 OTUs were specifically observed in blue, white and red light, respectively. The overall OTU diversity decreased with water depth, where deeper-water corals show the largest number of missing OTUs (Fig. S3). However, most OTUs were found at mid-depth, likely reflecting the higher number of coral samples collected from intermediate depth.

For downstream statistical analyses, we wanted to focus on the most abundant *Ostreobium* OTUs, and filtering out the low-abundance (<100 reads in total, and present in <3 samples) OTUs left us a final set of 27 abundant OTUs used in the downstream analyses. These are shown with red stars in Fig. 2A.

### Natural *Ostreobium* communities are strongly structured with depth

Principal coordinate analysis (PCoA) biplots based on abundances of *Ostreobium* OTUs in pre-treatment samples mainly grouped samples according to the water depth at which they were collected (Fig. 3). Shallow-water samples (red-yellow) separated from mid depth (green) and deeper-water (blue) samples along the PCo1 axis (with 35.1% of the variation explained by this component), while deeper-water samples separated from mid depth samples along PCo2 (Fig. 3). This result indicates that the *Ostreobium* community composition is vertically structured across the sampled depth gradient. More specifically, we were able to identify 8 abundant OTUs that drive most of the variation observed between depths in pre-treatment samples (grey bars in Fig. 4). OTUs 1, 2 and 27 are mainly found in shallow-water samples. While OTU 27 appears to be a shallow-water specialist not observed in mid- and deeper-water samples, OTUs 1 and 2 seem to have a more generalist nature, being present in pre-treatment samples from all depths, even though they are most common in shallow water. OTUs 17, 36 and 38 appear to be mid-water speciaists, and OTUs 16, 22, and 30 were predominantly found in deeper-water samples and barely observed in shallow-water samples. These OTUs are the same ones driving most of the structure observed in the pre-treatment biplot (Fig. 3).

**Figure 3.**
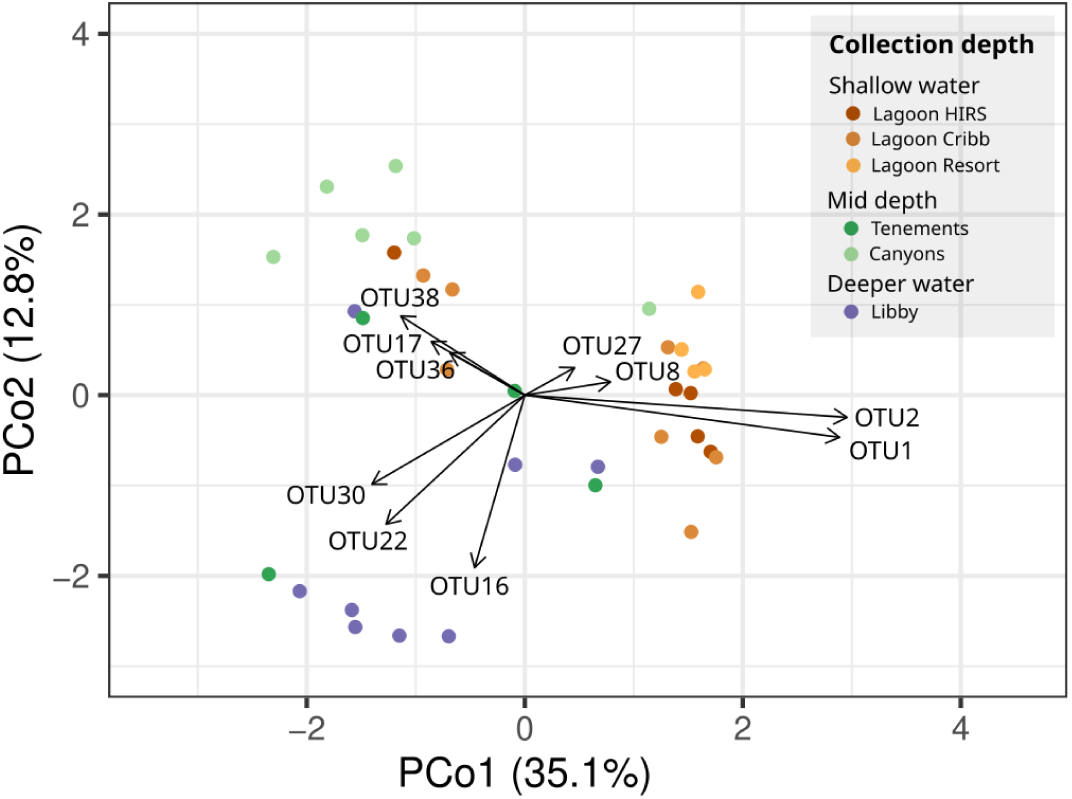
*Ostreobium* community structure in natural conditions. Two-dimensional representation (principal coordinates analysis) of the *Ostreobium* community structure, indicating the OTUs explaining most of the variability observed between samples collected from different depths.

**Figure 4.**
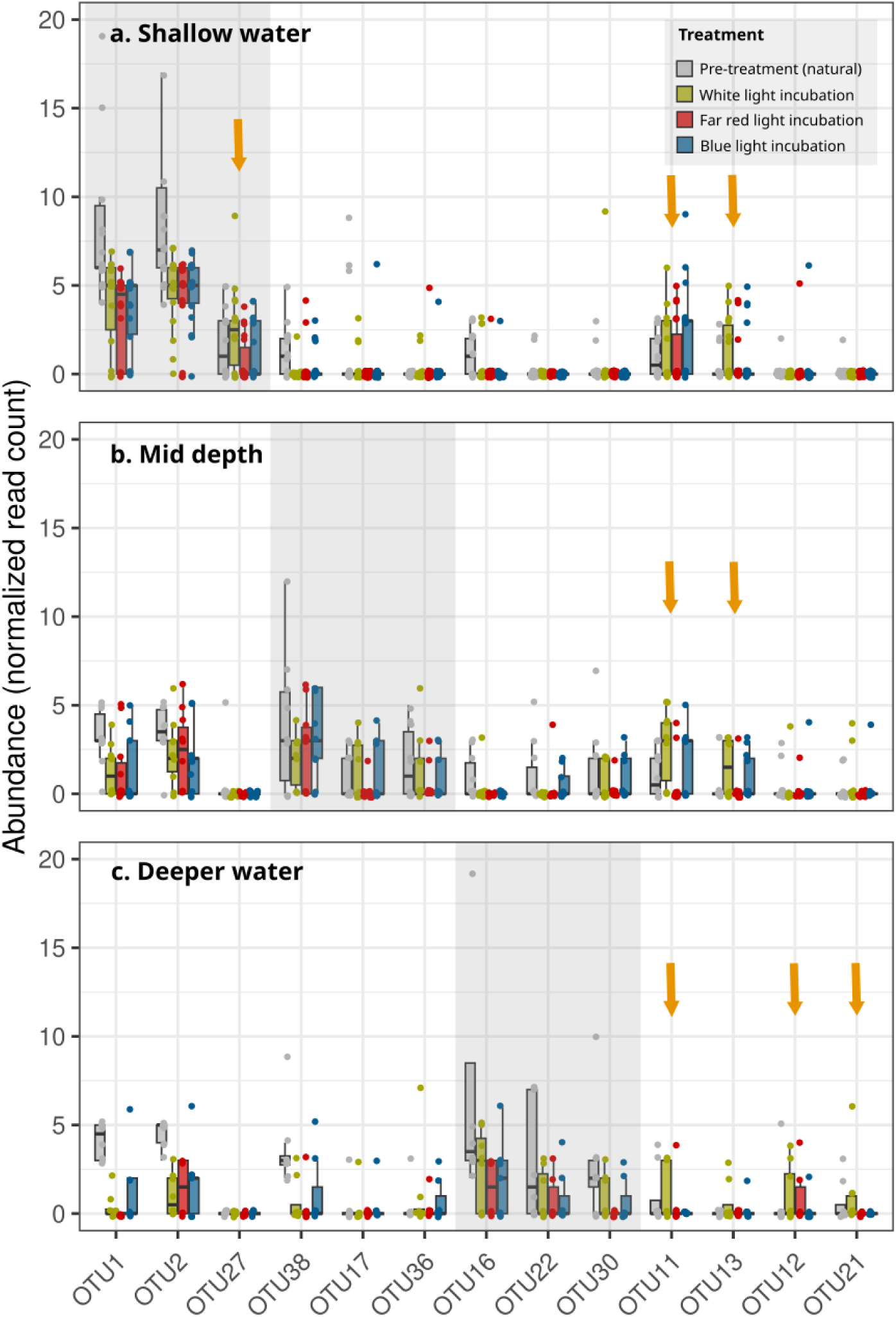
*Ostreobium* OTU profiling in natural conditions and light treatments. Boxplots showing the abundance of the 13 most abundant *Ostreobium* OTUs in different light treatments. Different plots contain information on samples collected from different water depths: **(a)** Shallow water, **(b)** Mid depth, **(c)** Deeper water. The grey background shading highlights 9 OTUs that stand out as the most prevalent OTUs at specific water depths before light treatment. Arrows highlight the five OTUs that show an increased abundance after light treatment.

### Light treatment identifies generalist and specialist OTUs

The overall response of OTUs to light treatment is shown graphically in Fig. 4 for a selection of OTUs, with counts and log-fold change statistics for a more extensive list of OTUs shown in Tables S1 and S2, respecively. These results all indicate that a range of responses can be observed across *Ostreobium* OTUs.

The most immediate observation is that the two OTUs that dominate in nature (1 & 2) reduced substantially in relative abundance following incubation in any type of light (Fig. 4). OTUs typical of shallow waters in nature (1, 2 & 24) reacted similarly to all light conditions. It is also clear that the OTUs that were more abundant in nature in mid-(38, 17 & 36) and deeper waters (16, 22 & 30) generally performed better in blue and white-light incubations than in far red (Fig. 4). Of the highly abundant OTUs shown in Fig. 4, only a handful increased in relative abundance following incubation (27, 11, 13, 12, 21; orange arrows), with rarer OTUs, most of which are not shown in Fig. 4, increasing in abundance more frequently (Table S2).

Of the five OTUs from Fig. 4 that were seen increasing, OTU27 was specifically found in shallow samples in nature and it performs best in white light (*LFC_white_*>5), OTU11 was found in nature at all depths and did particularly well in white light (*LFC_white_*>6), OTU13 was rare in nature at all depths but its abundance enhanced substantially in white light treatment (*LFC_white_*>7). OTUs 12 and 21 were rare in nature, but showed a positive response to white light incubation in samples originating from deeper water, with OTU 12 also increasing in far red light.

Roughly half of the 27 OTUs included in our analysis were considered generalists, performing similarly across most or all conditions (similar *LFC*s; Table S2). Ten OTUs were considered white light specialists (high SI), as well as two blue specialists (OTUs 24 & 26). Six OTUs were considered to thrive in far-red light based on their *LFC_far-red_* (4, 11, 12, 13, 28 & 35; Table S2), of which one (OTU 4) was considered a far-red specialist, as it performed much better in far-red incubation than in other incubations.

We provide more detailed report cards of the OTUs, including our interpretations of observed trends, in Supplementary Text 1.

### Pigment profiles reflect the photosynthetic light niche

In *Ostreobium* communities obtained from natural conditions (pre-incubation), chl*a* decreased with the depth of collection, with the highest value found for samples from shallow water and gradually declining towards deeper water (grey bars in Fig. 4a vs. 4b vs. 4c). No significant variation was observed for chl*b* for the different depths among the pre-incubation samples (grey bars in Fig. 4d vs. 4e vs. 4f).

Substantial differences in chlorophyll concentrations could be observed following incubation, with an overall trend towards lower concentrations in far-red light, higher concentrations in blue light, and intermediate concentrations in white light (Fig. 5).

**Figure 5.**
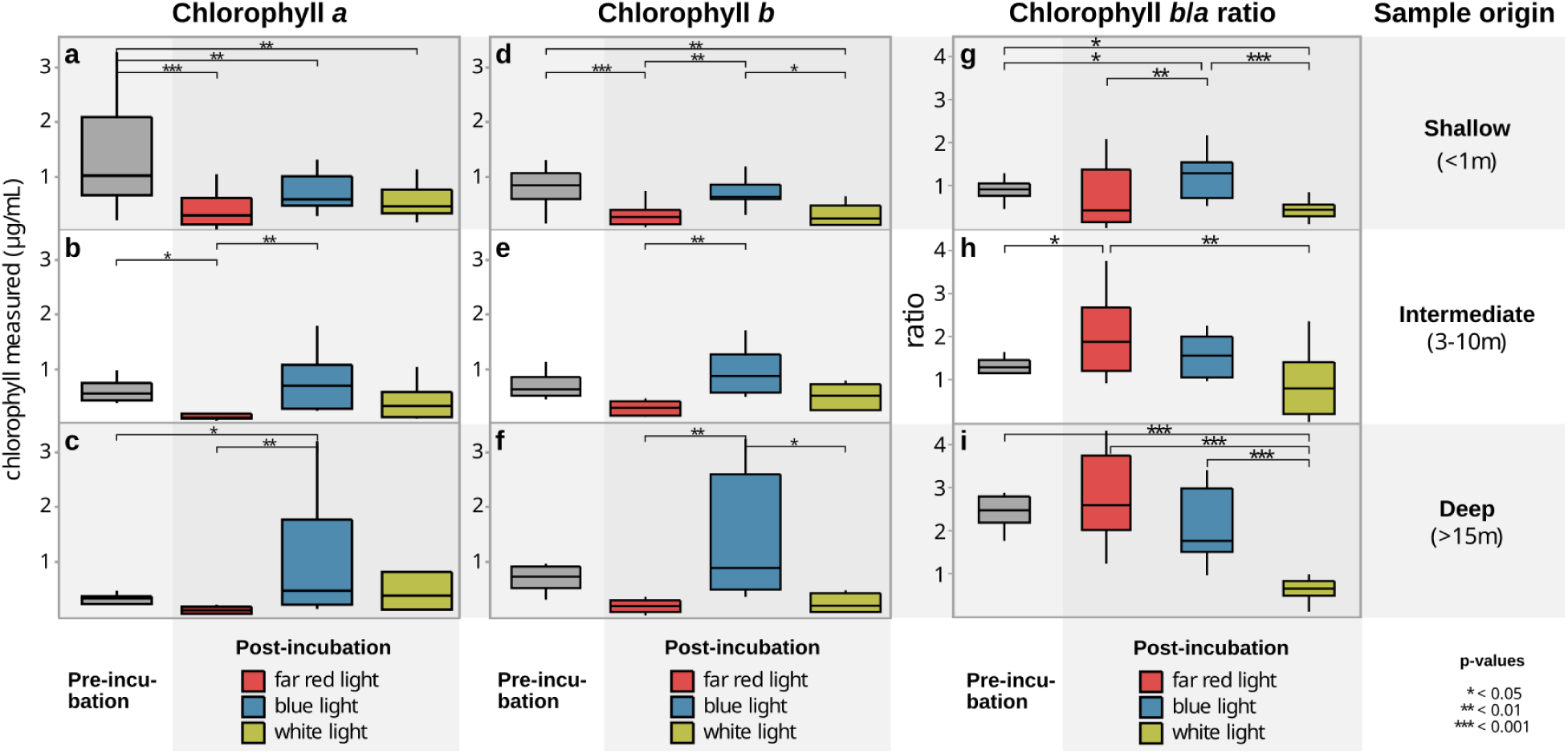
Concentration of chlorophyll *a* and chlorophyll *b* of *Ostreobium* collected from different depths and incubated in different light conditions. **(a,b,c)** Concentration of chlorophyll *a* for samples collected at shallow, intermediate and deep sites. **(d,e,f)** Concentration of chlorophyll *b* for samples collected at shallow, intermediate and deep sites. **(g,h,i)** Chlorophyll *b* to *a* ratios for samples collected at shallow, intermediate and deep sites. Samples collected in natural habitats (pre-incubation) are in grey, while colored bars respond to different light treatments.

With far red-light incubation, chl*a* and chl*b* were both substantially reduced compared to the pre-incubation concentrations, regardless of the original depth at which samples were collected (Fig 5). On average, the chl*a* value following incubation in far red light was higher in samples collected from shallow waters compared to other depths (compare red bars between Fig. 5a vs. 5b and 5c, p<0.05). There were no significant differences in chl*b* concentrations between samples from different depths following far-red light incubation (Figs 5b, 5d, 5f).

There was substantial across-sample variation in chlorophyll *a* and *b* concentrations following incubation in blue light, as indicated by the high variance indicated by the wide blue barplots (Fig. 5). In the majority of cases, the chlorophyll concentrations following blue light incubation did not differ significantly from pre-incubation samples, but they were often significantly higher than those seen after far-red incubation. After white light incubation, neither chl*a* nor chl*b* differed between samples originating from different depths. Chlorophyll *a* and *b* decreased marginally from the pre-incubation conditions following white light incubation, with significant reductions for samples originating from shallow water.

The chl*b*/chl*a* ratio is a statistic measuring the relative investment in light-harvesting antennae (which contain clh*b*, among other pigments) and reaction centers (which contain only chl*a*), and is often used to indicate dark acclimation or adaptation. In our data, the observed ratios in natural light conditions (pre-incubation) increased with depth (Fig. 5g vs. 5h vs. 5i), following the expected pattern of deeper (darker) sites having higher proportions of chl*b*/chl*a*.

The post-incubation samples under white light showed a stark reduction of the chl*b*/chl*a* ratio (Fig. 5g, h, i), which was significant compared to the pre-incubation state for shallow (<1m) and deeper samples (>15m). Samples collected at intermediate depths still showed this trend of reduction, albeit not statistically significant due to a wider spread of the post-incubation values. Samples incubated under blue light barely differed from the pre-incubation samples, except for a slight increase of the chl*b*/chl*a* ratio for samples originating from shallow habitats (Figure 5g).

The chl*b*/chl*a* ratio for samples incubated in far-red light showed a tendency to have more spread in values among replicates compared to natural light condition, but no consistent trend of increase or decrease could be observed (though values were significantly higher for samples from intermediate depths).

### Niche preferences are phylogenetically structured

*Ostreobium* lineage 4 is largely composed of specialists, shown in Fig. 6 by the low selectivity index across that entire lineage. Many species were classified as white light specialists, but OTU 24, which is sister to all remaining OTUs in lineage 4, is a blue light specialist. Species in lineage 4 generally show poor performance using far red light, though there are a few exceptions.

**Figure 6.**
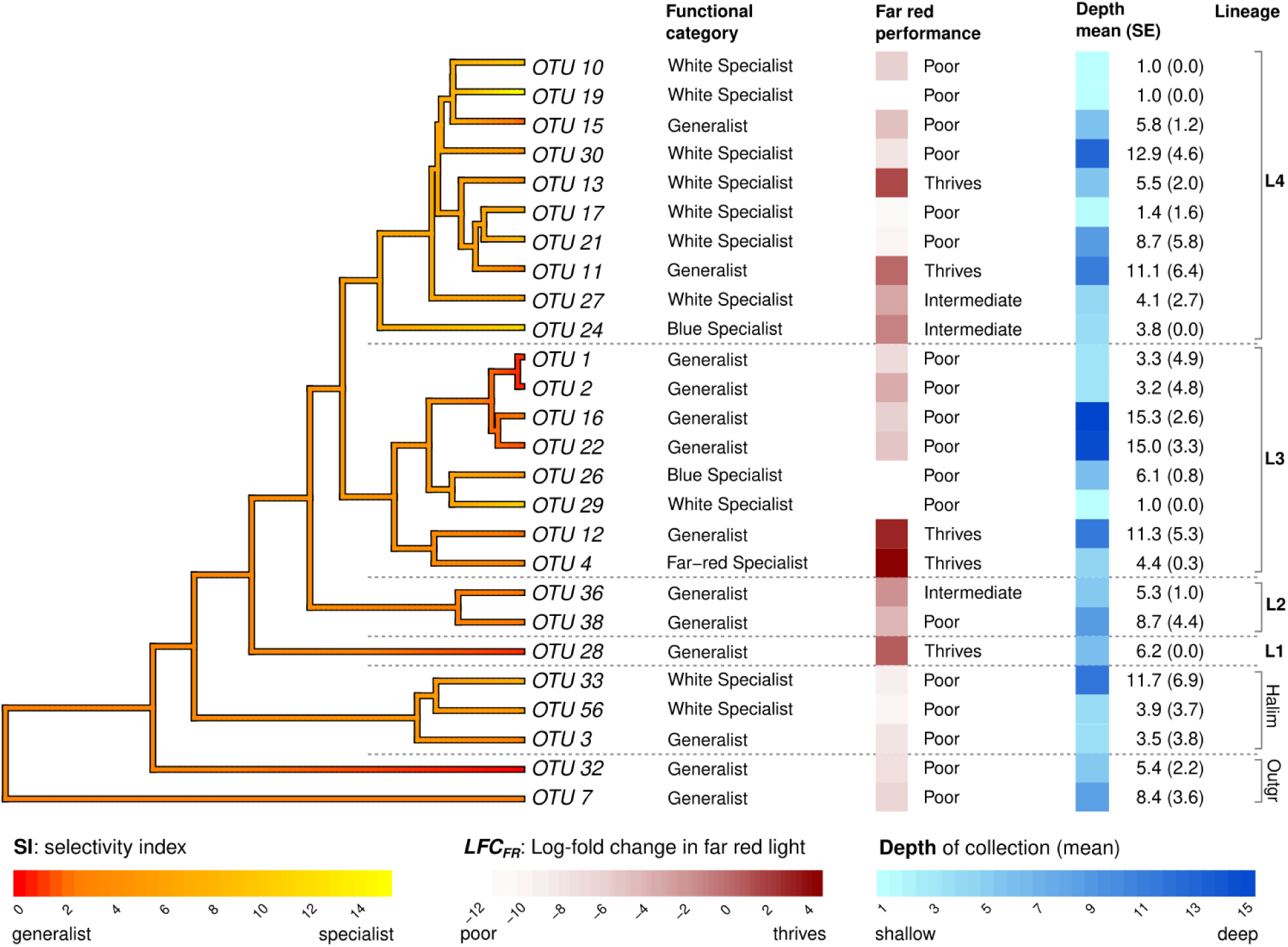
Phylogenetic structure of photobiological and microhabitat traits in *Ostreobium*.

Lineage 3 contains three main sublineages, one of generalists (OTUs 1, 2, 16, 22), one of specialists (OTUs 26, 29) and one that consists of two species with strong performance in far-red light, including OTU 4, which is considered a far-red specialist and OTU 12, which performed better in far red than in any other incubation but did not show extreme differences between incubations and is therefore classified as a generalist.

Lineage 2, for which we only have data for OTUs 36 & 38, both classified as generalists but showing an intermediate level of specialisation. OTU28 is the only representative from L1 and is classified as a generalist from mid-depth waters. It thrived in far-red light. Outside of *Ostreobium*, the two Halimedineae endolithic families, represented by OTUs 3 and 56, were shown to be generalists lacking far-red light photosynthetic capabilities.

## Discussion

Eco-physiological knowledge on *Ostreobium* has largely been derived from comparisons of culture strains (Koehne et al., 1999; Massé et al., 2020), which is naturally labor-intensive, leading to knowledge being rather anecdotal. Our work illustrates that gaining eco-physiological information can be scaled up substantially for difficult-to-culture organisms. Our design of experimental incubations, linked with high-throughput sequencing to characterise winner and loser lineages in a range of circumstances, has led to a leap in our knowledge of *Ostreobium* ecology and physiology across a large number of OTUs.

Our results show that the composition of the *Ostreobium* communities is structured along a depth gradient in nature, and is strongly altered by incubation in different light conditions. Our combined observations across natural habitat preferences and light incubations identify clear ecological generalists and specialists amongst *Ostreobium* OTUs, and assessment of chlorophylls show how communities alter their light harvesting machinery in nature and the incubations. Our observations help explain the patterns of diversity and distribution of *Ostreobium* across coral reef lightscapes and open perspectives for predicting relevant disturbance scenarios. Additionally, our results show clear phylogenetic conservatism of eco-physiological features and allow refining our knowledge of *Ostreobium* biodiversity and classification.

### Specialisation strategies

Our data show clear evidence for a range of different strategies among *Ostreobium* OTUs, from generalist species with similar responses to incubation in far-red, blue and white light, to specialists that perform particularly well in one specific light condition. The detailed report cards about performance of OTUs, along with our interpretations of the results (Supplementary Text 1) can serve as a reference to deepen the ecological interpretations of future environmental sequencing data, enriching the functional knowledge that can be obtained from otherwise descriptive surveys. We hope that future eco-physiological work on culture isolates of these OTUs and experimental work at the level of whole communities from other parts of the world and/or using other relevant incubation conditions can further deepen what we know about the ecology of different *Ostreobium* lineages. It is clear, though, that *Ostreobium* cannot longer be considered as ecologically homogeneous, and that interpretations require a more fine-grained view that distinguishes between OTUs.

The incubations showed OTU 4 to be a far-red specialist, with OTU 12 also performing better in far-red than in other incubations but not reaching the selection index threshold to be considered a far-red specialist, and several other OTUs also performed well in far-red incubation as part of their broader-spectrum light-harvesting strategy. Far red photosynthesis in *Ostreobium* has been shown to center around a red-shifted chlorophyll *a* configuration associated with light harvesting complex A1 (Lhca1), along with a highly uncommon coupling of this Lhca1 with photosystem II, allowing the splitting of water using the energy from far-red photons in an uphill energy transfer process (Koehne et al., 1999; Wilhelm & Jakob, 2006).

The blue light specialists identified in our experiment also performed reasonably well in white light, which contains a fraction of blue wavelengths, and this may contribute to explaining that these OTUs was mostly found in nature at mid-depths. A relatively large number of OTUs was classified as white light specialists. Philosophically, one could argue that there is no such thing as a white light specialist, since white light contains a wide span of wavelengths in the visible spectrum. Upon closer inspection of these white light specialists, we can distinguish between a subset that did well in exclusively blue wavelengths and may thus sustain itself in a narrow spectral band (e.g. OTUs 11, 13), while others did poorly in the blue incubation, indicating they require a wider range of wavelengths to achieve growth (e.g. OTUs 10, 19, 27). Furthermore, the OTUs classified as white specialists may just be the best responders to the relatively high light intensity present in this light treatment.

In addition to these specialists, we identified a range of generalist *Ostreobium* OTUs, including OTUs 1 & 2, with very high abundances in the Heron Island lagoon, and several other common OTUs of mid- and deeper waters (OTUs 38, 36, 16, 22). In our framework, generalists are defined as OTUs that are not very selective about the light condition they are exposed to. We used a selection index (SI) less than 3.16 as the threshold, meaning that the increase in relative abundance of the OTU in the condition where it grew best was less than a factor 2^3.16^ (=10) times the average of the change in the abundance of the other two conditions; in other words there is no huge change in the dynamics of the relative abundance of the OTU depending on the incubation. The same reasoning applied in case of negative growth (i.e. all incubations saw similar reductions). The threshold is to some extent an arbitrary choice to facilitate communication and align our work with the specialist-generalist theoretical framework, but the LFC and SI statistics can be interpreted directly for more nuanced interpretations along the specialist-generalist continuum.

It is important to note that our work is based on relative abundances, a constraint enforced by the nature of environmental sequencing, that requires some nuance in interpretations. First, an increase in relative abundance of an OTU does not automatically imply growth of that OTU. It just means the OTU did better than other OTUs. It may well be that all OTUs are reducing in absolute terms, but that this OTU is reducing less than others. Analogous reasoning applies to reductions in relative abundance, which can happen even for actively growing OTUs if many other OTUs are growing even more profusely. This complicates interpretations somewhat, because the performance of an OTU is measured in relation to whichever other OTUs happen to be present in a sample, and there will be a certain level of stochastic variation in this. Our results do facilitate a ranking in the level of specialisation of strains, and the fact that there is phylogenetic conservatism in spectral performance profiles enhances the biological credibility of these results. Furthermore, in related algae where growth in different light spectra has been investigated with culture strains, there are clear trends in growth rates and photosynthetic rates (Kang et al., 2020), and co-cultures reinforce the notion that light spectrum affects interspecies competition (Tan et al., 2020).

### *Ostreobium* in the spectral architecture of the reef

Our data provides clear evidence for particular *Ostreobium* OTUs dominating at different water depths. While this is a new observation for the Great Barrier Reef, similar results have been obtained in the Red Sea (Gutner-Hoch & Fine, 2011) and the Caribbean Sea (Gonzalez-Zapata et al., 2018; Rodríguez-Bermúdez et al., 2023), suggesting a global bathymetric niche partitioning of *Ostreobium* strains. While we draw this parallel across these distant locations, we are not suggesting that the same OTUs drive the depth gradients found in different oceans. Due to mismatches in marker genes and sequencing strategies between studies, global-scale OTU-level comparisons remain difficult at present (but see Giraldo-Vaca & Sánchez, 2024). What we can infer is that analogous generalist and specialist strategies are likely to exist among the *Ostreobium* OTUs found in different coral reefs, resulting in similarly structured depth gradients in *Ostreobium* community structure.

Physiological information relevant to the light environment on the reef can be gleaned from our chlorophyll measurements, which largely reflect light quantity, with some nuances related to spectral qualities. Natural communities had higher chl*b*/*a* ratios in deeper water, reflecting their larger light harvesting antennae (Koehne et al., 1999). Similarly, the lower-intensity blue light incubations, designed to resemble deeper-water light environments, had higher chl*b*/*a*.

The significant drop of chl*a* and *b* in the white light incubation for shallow-water communities is interpreted as a photoacclimation response to the increased light quantity (10 µmol m⁻² s⁻¹) in these incubations, favoring a reduction of chlorophylls to enhance energy efficiency and reduce photoinhibition. In line with this, the substantial drop in the chl*b*/*a* ratio signals a reduction of the photosynthetic antenna size. The stability of chl*a* and *b* for mid- and deeper-water communities incubated in white light suggests they rely more on remodelling their light harvesting complexes while maintaining overall chlorophyll concentrations. These interpretations at the community level are complicated somewhat by the underlying changes in relative abundances of OTUs, so the level to which we see acclimation of OTUs versus changes in their relative abundances is hard to evaluate.

The drop in both chl*a* and *b* in far-red incubations is probably best explained as a reduction of overall biomass in these conditions inhospitable to the majority of OTUs. Theoretically, one may expect far-red light incubation to favor a higher relative amount of chl*a*, since the current paradigm suggests that far-red photosynthesis is facilitated by red-shifted configurations of chlorophyll *a* (Fork & Larkum, 1989; Koehne et al., 1999; Wilhelm & Jakob, 2006). However, counter to this theoretical expectation, we observed higher chl*b*/*a* ratios in far-red light than in white light incubations. Perhaps this is a response of the predominant strains who do not use far-red wavelengths and would therefore perceive this incubation as very low-light, which would stimulate a shift towards chl*b*.

Our results show that the spectral niche framework is useful in understanding the biology of *Ostreobium* in the sense that major variation in the spectral preferences was observed among OTUs. However, we also show that this framework may not be the best predictor of *Ostreobium* distributions in nature, since our experiments suggest that many *Ostreobium* OTUs may be operating away from their optimal spectral environment. For instance, in deep water, where one may expect blue specialists, the two most abundant OTUs (16, 22) are generalist doing well in all incubations. The blue specialists (OTUs 24, 26), on the other hand, were not very abundant at the deepest sites. White specialists were found from the shallowest to the deepest sites, so do not appear to have specific preferred sites in nature.

These results indicate that the realised spectral niche of OTUs in nature is much narrower than their fundamental spectral niches. Mismatches between fundamental and realised niches are fairly common in nature. For instance, salt-tolerant plants often grow better in freshwater when in isolation, but lose out in such environments in nature due to competitive exclusion, while they win the competition in higher-salinity habitats (Crain et al., 2004). In water-deprived habitats, the desert cactus *Opuntia* grows better in isolation when given water, but in artificially watered desert plots, cacti lose out to grasses (Briones et al., 1998).

For *Ostreobium*, other aspects of the abiotic niche may contribute to restricting realised niches, including skeletal density properties and nutrient availability (Fordyce et al., 2021). However, we think that it is primarily biotic interactions, more specifically competition amongst phototrophic organisms, that play a major role in determining which *Ostreobium* OTUs dominate in particular environments in nature. For instance, the OTUs most performant in far-red incubations were not found primarily in shallow waters, where one might expect such species to thrive, but in mid-depth (OTU 4) and even deeper (OTU 12) waters. In shallow-water corals, they face competition from generalist and white-specialist *Ostreobium* OTUs, along with a range of other photobionts that have more advanced mechanisms to capture far-red light, including cyanobacteria with chlorophylls *d* or *f* (Chen, 2019). The uphill energy transfer required for far-red photosynthesis in *Ostreobium* comes with a significant reduction in maximum quantum yield (Wilhelm & Jakob, 2006), a trade-off that may prevent such strains from dominating shallow waters.

Similarly, some white light specialists appear to be restricted to deeper blue waters, not because blue is their favored wavelength, but because this is a niche where they do not get outcompeted by shallow-water generalist OTUs. The OTUs showing good performance in blue wavelengths, on the other hand, were not deep-water specialists, but found mostly in mid- to shallow-water habitats in nature, where blue wavelengths are common as well, but often used at least to some extent by the Symbiodiniaceae forming the equivalent of a canopy in the coral tissue above.

All in all, the results show that in nature, distributions do not always follow expectations based on spectral preferences. This signals that *Ostreobium* likely takes advantage of the diversity of wavelengths that are present in those natural environments. Shallow water coral skeletons do not exclusively receive far-red light, but also small amounts of other wavelengths that *Ostreobium* could take advantage of. Furthermore, light conditions inside the coral will undergo fluctuations depending on tides, which will be particularly noticeable in shallow water corals, and lineages that can utilise a wider range of light spectra will benefit. In this context, it’s important to note that our work focused mostly on blue and far-red specialisation. Work on some *Ostreobium* culture strains shows that accessory pigments including siphonein and (likely) siphonoxanthin are also present (Fork & Larkum, 1989; Schlichter et al., 2008; Wilhelm & Jakob, 2006). These can facilitate the use of green wavelengths to enhance photosynthesis, which Symbiodiniaceae do not deplete as much as blue and red. The relevance of these wavelengths to the photosynthesis of different *Ostreobium* strains remains an open question.

### Ecological implications and coral bleaching

The roles of *Ostreobium* in the coral holobiont have been discussed to some extent, and the question of how this alga, which may provide an alternative source of energy during coral bleaching events, is an important and urgent one to answer. Our results highlight that there will be no one single answer, but rather that *Ostreobium* is a physiologically heterogeneous group, so understanding its ecological roles and responses to environmental disturbances will require a more fine-grained approach that embraces the diversity within.

In natural circumstances, the coral skeletons are dominated by a handful of very abundant OTUs, with different ones dominating at different depths. Getting more detailed ecological information for these OTUs is paramount, since they are so crucial on the reef. Our work shows that these abundant taxa are nearly all generalists when it comes to their preferences in light spectra, having similar performance across most or all incubations.

One of the most striking ecological observations from our study is that nearly all the common OTUs in nature drastically reduced in relative abundance in our incubations, regardless of which type of light they were exposed to. This includes OTUs 1 and 2, the overlords of the shallow reef with abundances towering over those of all other OTUs. They remain relatively common even after incubation, but this is largely due to their strong starting position, as their relative abundance dropped by 6- to 90-fold depending on the incubation type. This result is counter-intuitive. They are classified as generalists based on photobiological parameters, clearly win the competition in nature, yet are struggling when conditions change while other OTUs are thriving. Explanations for this likely have to be found in other dimensions of the abiotic niche and/or biological interactions. An intriguing perspective, albeit purely speculative at this stage, is that these strains may depend more strongly on biotic interactions in nature, potentially cross-feeding with particular bacteria or the coral itself, for instance in the supply of vitamin B12 and nitrogen compounds (Iha et al., 2021).

In changing environments, specialist species have been shown to be declining faster than generalist species (e.g., Olden et al., 2004), leading to functional homogenization of communities. Since climate change increases the frequency and severity of coral bleaching, one may argue that generalist *Ostreobium* species might be favoured as well, particularly in the skeletons of shallow-water corals, where the light environment, which usually consists of low light enriched in far-red wavelengths, would change more regularly to high-intensity white light as corals bleach more often.

Following this logic, the generalist strains that are currently dominant in nature would remain firmly in charge. However, this reasoning is strongly challenged by our experimental incubations in white light, which to some extent mimic the light environment seen in bleached coral (Galindo-Martínez et al., 2022), and which appear detrimental to these dominant OTUs. Even though we consider these OTUs generalists, our definition is limited to their light spectral range, and it is quite possible that these lineages do not fare well in the higher-light environments of our white-light incubations, which would extend to bleached coral.

Particularly the white specialists identified in our work are the primary targets to watch as potential “winners” in the era of heightened coral bleaching. In our opinion, it will be particularly the species tolerant of the high light levels present in coral skeletons during bleaching that will differentiate the winners from the losers. Isolating culture strains of the white light specialists to gain better understanding of their physiology and comparing it to that of the dominant generalist OTUs should be a priority to investigate their physiology and metabolism, and facilitate predictions about their dynamics during coral bleaching.

Of course, any such experiments, including our own, are highly simplified, and several other factors change alongside the light environment in coral bleaching events. So, alongside this type of work, it will be highly beneficial to track the biological responses of these OTUs along with those of the host and its microbiome in experimentally or naturally bleached corals.

Information based on comparison of photosystem (*psb*A) transcript abundances in bleached and unbleached corals has already indicated different responses among *Ostreobium* OTUs in bleached coral (Iha et al., 2021), and based on our analysis of these *psb*A sequences, those with enhanced activity are mainly in L4, providing anecdotal information that this group in which many white light specialists are found, may indeed increase in importance during bleaching events.

### Evolutionary and biodiversity discoveries

There is strong phylogenetic conservatism of niche preferences, with generalists and specialists mostly falling out in different lineages. Particularly the subclade of L3 consisting of OTUs 1, 2, 16 and 22 consist entirely of functionally similar generalists. This strategy has allowed it to dominate the reef, with OTUs 1 & 2 as the most abundant OTUs in shallow water, and OTUs 16 & 22 found as dominant species in deeper water. Lineage 4 is home to the blue and white specialists. This lineage appears to be a very diverse one, with many lineages sprouting from a poorly resolved backbone (Fig. 2), potentially indicating a rapid radiation, although the short amplicons and conservative marker used here may also contribute to this pattern.

Far-red photosynthetic capabilities are not universal, but seem widespread across the *Ostreobium* tree, including several of the main lineages: L1 (OTU 28), L3 (the OTU 4+12 clade) and two unrelated OTUs in L4 (11 and 13). Comparison between *Ostreobium* culture strains (Koehne et al. 1999) identified a unique light harvesting complex protein (Lhca1) in strains exhibiting far-red absorption, and this modified protein may be involved in the capacity for *Ostreobium* to extend the absorption of its antennae into far red. The *Ostreobium* genome of strain SAG6.99 (in L4; see Fig. 2) showed an extensive set of light harvesting complex proteins (Iha et al 2021), including expansion of the LHCA subfamily, but it is not clear how precisely the various LHC proteins and pigment combinations contribute to *Ostreobium*’s photosynthetic capabilities in different light spectra. However, based on the widespread occurrence of far-red capable lineages across the *Ostreobium* tree, we hypothesise that far-red photosynthesis may have its origin at the base of the tree, and was subsequently in several lineages. The whole genome duplication that has recently been inferred to have happened at the origin of the Ostreobineae (Hossen et al., 2026) may have played a role in the gain of far red photosynthetic capabilities by duplication and neofunctionalisation of LHC genes (Ohno, 1970).

The OTUs encountered in our coral skeletal samples show that *Porites* from the Great Barrier Reef is home to a diverse range of *Ostreobium* lineages, as well as some OTUs from other bryopsidalean suborders including the endolithic families mentioned in previous biodiversity surveys (Marcelino & Verbruggen, 2016; Sauvage et al., 2016). We found that OTU 101 likely represents an additional deep-branching lineage (L5), and with these lineages estimated to be ca. 500 million years old (Marcelino & Verbruggen, 2016), this discovery adds substantially to known phylodiversity. More broadly, our results clearly confirm the substantial unrecognised biodiversity within *Ostreobium*. Only a handful of *Ostreobium* species have been described, yet evidence from culture studies and environmental sequencing work such as ours shows that this is a drastic underestimate (refs), likely due to a combination of rock-dwelling algae being understudied by taxonomists and the simple structure of the genus not leaving much room for morphological differentiation (e.g., Díaz-Tapia et al., 2020; Verbruggen et al., 2009), leading to an accumulation of cryptic species over the lineage’s long evolutionary history.

The many alternative classifications that have been applied to *Ostreobium* are a particularly hard nut to crack. This has been largely due to different studies using different markers: 16S rRNA (del Campo et al., 2017), *tuf*A (Marcelino & Verbruggen, 2016; Sauvage et al., 2016), and *rbc*L (Gonzalez-Zapata et al., 2018; Gutner-Hoch & Fine, 2011; Massé et al., 2020). The lineage classifications based on these different markers remain difficult to match up due to there not being much overlap between markers, as much of the biodiversity surveys have been done via environmental metabarcoding. Our work advances that field by reconciling the main classifications used for the *tuf*A and 16s rRNA markers, thanks to the availability of paired *tuf*A and 16S rRNA data of chloroplast genomes (Alesmail et al., 2023; del Campo et al., 2017; Marcelino et al., 2016; Pasella et al., 2022; Verbruggen et al., 2017) and amplicon sequences of culture strains isolated by Thomas Sauvage (Sauvage et al., 2016; and additional sequences published on Genbank). Further work will be needed to also reconcile the taxonomic framework developed based on the *rbc*L marker, for which less comparative data is available. Lastly, the classification framework needs to be refined beyond just the main deep-branching lineages, as our work clearly shows sublineages with particular physiological features (e.g. the OTU1+2+22 generalist group; the OTU4+12 far red specialists), and naming these would greatly facilitate communication.

## Acknowledgements

This work was funded by the Australian Research Council (DP200101613 to HV) and the Holsworth Wildlife Research Endowment (to MMP). We thank Thomas Sauvage for publishing 16S rRNA sequences of a selection of his strains on Genbank, which aided the comparison of taxonomic schemes. We are grateful to the staff of the Heron Island Research Station for facilitating field work.

## Author contributions

MMP and HV conceptualised the study. MMP led the execution of the experiment, with significant contributions from FR during field work and AS and FP in aspects of molecular work. MMP, MP and HV performed data analyses. HV drafted the manuscript, with contributions of original writing from MMP and MP. Edits were contributed by all authors.

---

**Figure S1:**
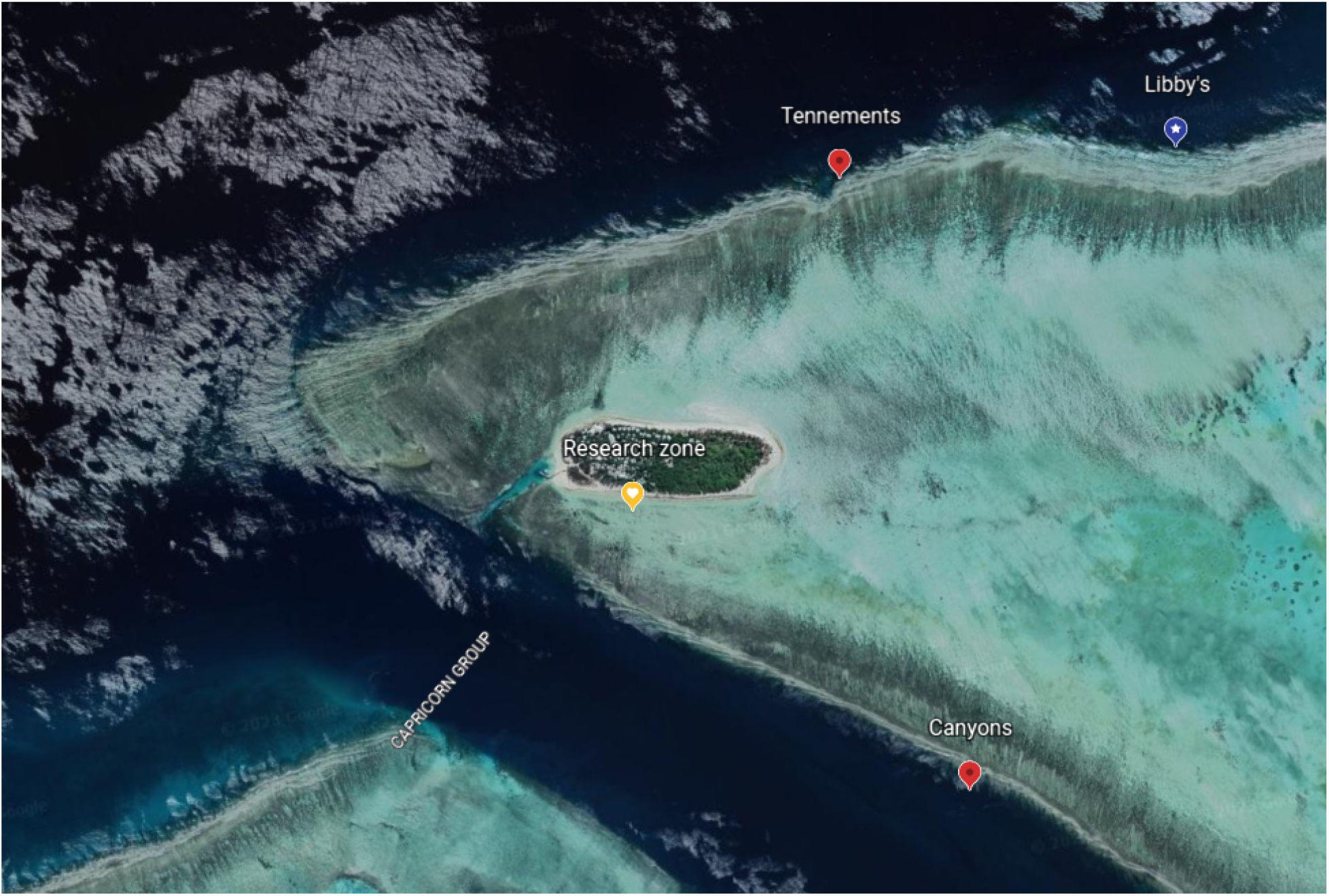
Sampling locations of the study on Heron Island, Great Barrier Reef, Australia.

**Figure S2.**
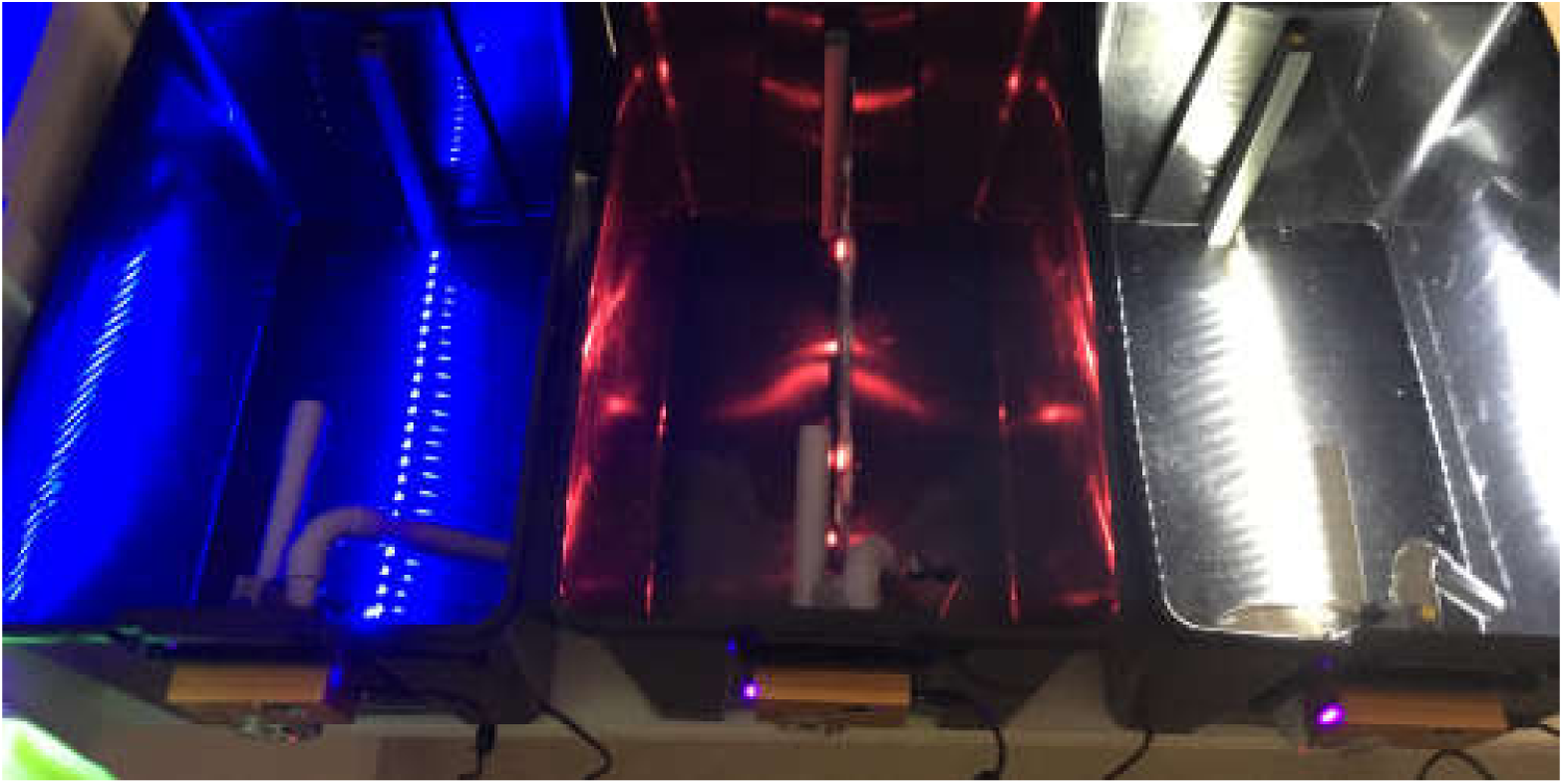
Photograph of the black boxes set up prior to incubating the coral fragments.

**Figure S3.**
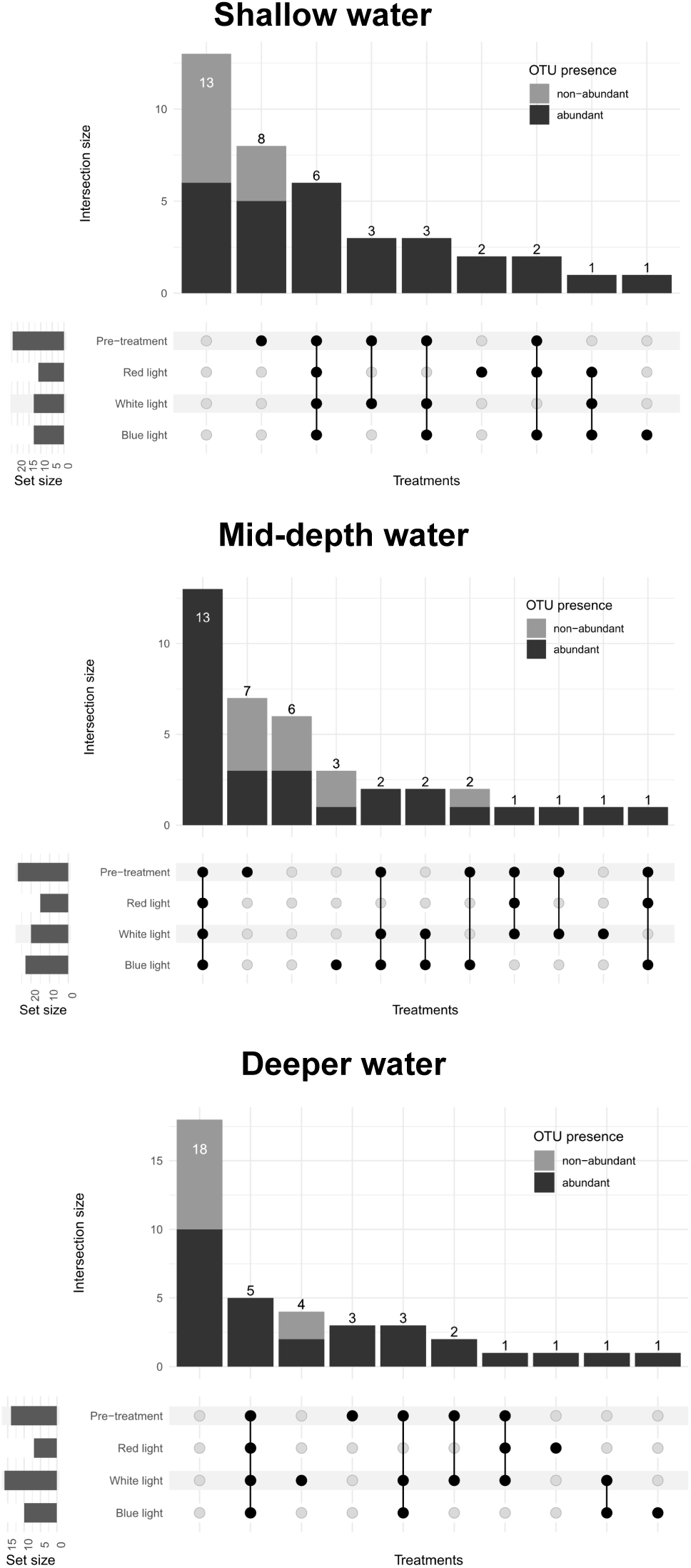
Depth-specific presence/absence upset plots. The plots represent the presence/absence of 39 *Ostreobium* OTUs throughout light treatments. Samples from different water depths were plotted separately: shallow (0-1 m), intermediate (1-9 m), deep (9-17.5 m). See main Figure 4 for analysis using samples from all depths.

**Table S1.** Total OTU abundances. The table illustrates the total OTU counts before and after light treatment. The different colors highlight the specialist OTUs that are only found in white, far-red, or blue light conditions. Furthermore, generalist OTUs are separated by OTUs that are only detected before light exposure.

| OTU | Pre-treat. | Light treatment |  |  |  |
| --- | --- | --- | --- | --- | --- |
|  |  | White | Far-red | Blue |  |
| OTU_25 | 0 | 670 | 0 | 0 | Specialists |
| OTU_29 | 2905 | 117 | 0 | 0 |  |
| OTU_101 | 0 | 15 | 0 | 0 |  |
| OTU_23 * | 0 | 0 | 191 | 0 |  |
| OTU_7 * | 115 | 0 | 0 | 7 |  |
| OTU_44 | 0 | 0 | 0 | 40 |  |
| OTU_51 | 0 | 0 | 0 | 53 |  |
| OTU_109 * | 0 | 0 | 0 | 120 |  |
| OTU_1 | 1275371 | 48417 | 21714 | 17352 | Generalists |
| OTU_2 | 1665462 | 61388 | 28899 | 22400 |  |
| OTU_3 * | 3011 | 0 | 12 | 7 |  |
| OTU_4 | 3356 | 284 | 1080 | 492 |  |
| OTU_10 | 46 | 235 | 0 | 153 |  |
| OTU_11 | 1427 | 23609 | 15138 | 69767 |  |
| OTU_12 | 1801 | 2124 | 2197 | 1371 |  |
| OTU_13 | 888 | 7249 | 420 | 6136 |  |
| OTU_15 | 681 | 1131 | 126 | 1363 |  |
| OTU_16 | 117659 | 20184 | 283 | 4647 |  |
| OTU_17 | 18010 | 3750 | 25 | 9877 |  |
| OTU_19 | 3815 | 33420 | 0 | 62 |  |
| OTU_20 | 23 | 26 | 0 | 18 |  |
| OTU_21 | 913 | 12229 | 0 | 6294 |  |
| OTU_22 | 14301 | 435 | 197 | 1196 |  |
| OTU_24 | 985 | 2272 | 160 | 2159 |  |
| OTU_26 | 709 | 99 | 0 | 71 |  |
| OTU_27 | 7339 | 5502 | 625 | 1558 |  |
| OTU_28 | 52 | 344 | 34 | 1023 |  |
| OTU_30 | 26726 | 2981 | 15 | 1932 |  |
| OTU_33 * | 396 | 1681 | 0 | 79 |  |
| OTU_35 | 56 | 24 | 197 | 0 |  |
| OTU_36 | 7595 | 14743 | 200 | 2218 |  |
| OTU_38 | 100213 | 1600 | 3645 | 18616 |  |
| OTU_56 * | 3698 | 799 | 0 | 78 |  |
| OTU_32 * | 186 | 0 | 0 | 0 | Only in Pre-treatment |
| OTU_43 | 64 | 0 | 0 | 0 |  |
| OTU_54 * | 14 | 0 | 0 | 0 |  |
| OTU_55 * | 47 | 0 | 0 | 0 |  |
| OTU_86 | 49 | 0 | 0 | 0 |  |
| OTU_99 * | 7 | 0 | 0 | 0 |  |
| OTU_110 | 2 | 0 | 0 | 0 |  |

\* OTUs that fall outside the Ostreobineae suborder

**Table S2.** Classification of *Ostreobium* OTUs into generalist or specialist categories based on log-fold change values in their relative abundances following light incubation.

|  | LFC (Blue) |  | LFC (Far red) |  | LFC (White) |  | maximum | mean of | selection | Category | Far red |
| --- | --- | --- | --- | --- | --- | --- | --- | --- | --- | --- | --- |
|  | mean | SE | mean | SE | mean | SE | LFC | others | index |  | performance |
| OTU1 | -5.075 | 1.143 | -6.542 | 1.113 | -5.468 | 1.133 | -5.075 | -6.005 | 0.929 | Generalist | Poor |
| OTU2 | -2.632 | 0.745 | -3.472 | 1.126 | -2.933 | 0.802 | -2.632 | -3.203 | 0.571 | Generalist | Poor |
| OTU3 | -10.125 | 1.295 | -8.068 | 3.935 | -11.951 | 0.784 | -8.068 | -11.038 | 2.969 | Generalist | Poor |
| OTU4 | 0.995 | 15.065 | 4.264 | 6.291 | 0.762 | 7.976 | 4.264 | 0.878 | 3.386 | Far-red Specialist | Thrives |
| OTU7 | -3.409 | 3.148 | -6.104 | 2.345 | -6.280 | 1.667 | -3.409 | -6.192 | 2.782 | Generalist | Poor |
| OTU10 | 0.054 | 8.170 | -5.853 | 2.263 | 6.509 | 0.643 | 6.509 | -2.900 | 9.409 | White Specialist | Poor |
| OTU11 | 3.759 | 2.085 | 0.252 | 2.386 | 4.574 | 1.947 | 4.574 | 2.006 | 2.568 | Generalist | Thrives |
| OTU12 | 0.139 | 4.333 | 3.043 | 4.059 | 1.966 | 3.920 | 3.043 | 1.053 | 1.991 | Generalist | Thrives |
| OTU13 | 4.463 | 2.812 | 1.479 | 3.622 | 7.245 | 2.045 | 7.245 | 2.971 | 4.274 | White Specialist | Thrives |
| OTU15 | -1.227 | 5.619 | -4.714 | 5.943 | -1.397 | 5.702 | -1.227 | -3.055 | 1.829 | Generalist | Poor |
| OTU16 | -5.542 | 1.177 | -5.929 | 1.275 | -3.440 | 1.459 | -3.440 | -5.735 | 2.295 | Generalist | Poor |
| OTU17 | -2.757 | 3.543 | -10.986 | 1.963 | -0.124 | 3.049 | -0.124 | -6.871 | 6.747 | White Specialist | Poor |
| OTU19 | -5.048 | 8.819 | -13.867 | 0.036 | 6.214 | 4.430 | 6.214 | -9.458 | 15.671 | White Specialist | Poor |
| OTU21 | -5.079 | 4.151 | -9.595 | 0.237 | 1.493 | 3.497 | 1.493 | -7.337 | 8.830 | White Specialist | Poor |
| OTU22 | -4.738 | 2.009 | -5.214 | 2.651 | -7.340 | 2.606 | -4.738 | -6.277 | 1.539 | Generalist | Poor |
| OTU24 | 15.133 | 1.236 | -0.975 | NA | 8.012 | 7.449 | 15.133 | 3.518 | 11.615 | Blue Specialist | Intermediate |
| OTU26 | -0.799 | 3.963 | -11.541 | 1.052 | -1.202 | 4.608 | -0.799 | -6.372 | 5.573 | Blue Specialist | Poor |
| OTU27 | -1.217 | 2.632 | -2.890 | 2.701 | 4.074 | 1.976 | 4.074 | -2.053 | 6.127 | White Specialist | Intermediate |
| OTU28 | 2.185 | NA | 0.755 | NA | 2.017 | NA | 2.185 | 1.386 | 0.800 | Generalist | Thrives |
| OTU29 | -13.448 | 0.076 | -13.448 | 0.076 | -2.335 | 7.968 | -2.335 | -13.448 | 11.113 | White Specialist | Poor |
| OTU30 | -4.190 | 1.992 | -7.721 | 1.086 | -0.881 | 2.341 | -0.881 | -5.956 | 5.075 | White Specialist | Poor |
| OTU32 | -7.435 | 1.587 | -7.435 | 1.587 | -7.435 | 1.587 | -7.435 | -7.435 | 0.000 | Generalist | Poor |
| OTU33 | -1.453 | 7.342 | -8.787 | 0.631 | 2.334 | 5.751 | 2.334 | -5.120 | 7.454 | White Specialist | Poor |
| OTU35 | NA | NA | 3.461 | NA | -1.794 | NA | NA | NA | NA |  | Thrives |
| OTU36 | 2.208 | 3.438 | -1.787 | 3.948 | 2.475 | 2.890 | 2.475 | 0.210 | 2.265 | Generalist | Intermediate |
| OTU38 | -2.435 | 1.062 | -4.095 | 1.057 | -5.290 | 0.840 | -2.435 | -4.693 | 2.258 | Generalist | Poor |
| OTU56 | -7.649 | 1.889 | -9.498 | 0.427 | -3.101 | 2.558 | -3.101 | -8.574 | 5.473 | White Specialist | Poor |

## Supplementary text 1: Report cards of noteworthy *Ostreobium* OTUs

OTUs 1 and 2: These are dominant generalists, particularly in shallow water but also in mid- and deeper water. They do fine in incubations in any light condition, yet lose ground to other OTUs. These strains do not do well at all in incubations in far-red light, with their relative abundance dropping by 98.6% (OTU 1, *LFC_far-red_*=-6.177) and 85% (OTU 2, *LFC_far-red_*=-2.736). This is surprising, as the habitat in which they are found (and where they thrive) is heavily enriched in far-red light.

OTU 27: A lower-abundance OTU found near-exclusively in shallow water, which performs better in incubations than the dominant OTUs 1 & 2, increasing its relative abundance, particularly in white light, and it is considered a white light specialist. It does not perform well in far-red light, nor in blue light, indicating it needs a broader spectrum of wavelenghts to do well.

OTUs 17 and 36: These are mid depth specialists. They are fairly common in nature at mid depth only. They declined in incubations, but the samples collected from mid depths did relatively well in both white and blue light. They do very poorly in far red light, which indicates they may not possess the capability for uphill energy transport, and is in line with them only being observed in waters. These are classified as generalists.

OTU 38: This is a common species in nature, particularly dominant at mid depths, but also present in non-negligible quantities in shallow and deeper water. Like the other mid-water OTUs 17 and 37, it maintained its abundance fairly well in blue and white light. In contrast to OTUs 17 and 36, it had similar performance in far red light than in white and blue, reinforcing its generalist classification.

OTUs 16, 22 and 30: These are the most common species in deeper water. They decline in incubations, and are considered generalists since they show similar performance in most light conditions. The performance in FR is poor; they likely are incapable of uphill energy transport and are restricted to deeper water.

OTUs 11 and 13: Relatively rare (OTU 11) to very rare (OTU 13) in natural circumstances, but clear winners in incubations. They performed best in white light and were classified as white light specialists, but also perform well in blue light, which is unsurprising given that blue wavelengths represents a good fraction of white light. Both these strains thrived in far-red incubations, though much less so than in white and blue light.

OTUs 12 and 21: Only present in meaningful amounts in deeper water. They are both winners in our incubations, and both do well particularly in white light. They have different profiles in their spectral preferences though, with OTU 21 having poor performance in far-red, while OTU 12 thrives in this wavelength, which is contradictory to the OTU being mainly found in deep water where this wavelength would be largely filtered out by the water column.

OTU 24: This is among the rare OTUs in nature, but which does well in incubations, to the point that on several occasions, we observed it only following the incubation but not in the pre-treatment library. It occurs exclusively in mid and deeper-water sites. It is listed here mainly because it is considered a blue light specialist, though it has outstanding performance in white light too. It is classified as resilient to far-red, but with an LFC of -0.975 and a sparsity of data, it is probably best to consider this as an OTU that does not possess uphill energy transport.

OTU 4: As the previous OTU, this is among the rarer ones in nature, but does very well in incubations, and similarly has been observed in some samples only following incubation. It becomes particularly abundant in far red-light incubations (22x enhancement), and is therefore classified as a far red specialist likely to possess uphill energy transfer.

